# Mechanical stimuli activate gene expression via a cell envelope stress sensing pathway

**DOI:** 10.1101/2022.09.25.509347

**Authors:** Christine E. Harper, Wenyao Zhang, Jung-Ho Shin, Ellen van Wijngaarden, Emily Chou, Junsung Lee, Zhaohong Wang, Tobias Dörr, Peng Chen, Christopher J. Hernandez

## Abstract

In tissues with mechanical function, the regulation of remodeling and repair processes is often controlled by mechanosensitive mechanisms; damage to the tissue structure is detected by changes in mechanical stress and strain, stimulating matrix synthesis and repair. While this mechanoregulatory feedback process is well recognized in animals and plants, it is not known whether such a process occurs in bacteria. In *Vibrio cholerae*, antibiotic-induced damage to the load-bearing cell wall promotes increased signaling by the two-component system VxrAB, which stimulates cell wall synthesis. Here we show that changes in mechanical stress and strain within the cell envelope are sufficient to stimulate VxrAB signaling in the absence of antibiotics. We applied mechanical forces to individual bacteria using three distinct loading modalities: extrusion loading within a microfluidic device, compression, and hydrostatic pressure. In all three cases, VxrAB signaling, as indicated by a fluorescent protein reporter, was increased in cells submitted to greater magnitudes of mechanical loading, hence diverse forms of mechanical stimuli activate VxrAB signaling. Mechanosensitivity of VxrAB signaling was lost following removal of the VxrAB stimulating endopeptidase ShyA, suggesting that VxrAB may not be directly sensing mechanical forces, but instead relies on other factors including lytic enzymes in the periplasmic space. Our findings suggest that mechanical signals play an important role in regulating cell wall homeostasis in bacteria.

**Significance Statement:** Biological materials with mechanical function (bones, muscle, etc.) are often maintained through mechanosensitive mechanisms, in which damage-induced reductions in stiffness stimulate remodeling and repair processes that restore mechanical function. Here we show that a similar process can occur in bacteria. We find that mechanical stresses in the bacterial cell envelope (the primary load-bearing structure in bacteria) regulate signaling of a two-component system involved in cell wall synthesis. These findings suggest that the mechanical stress state within the cell envelope can contribute to cell wall homeostasis. Furthermore, these findings demonstrate the potential to use mechanical stimuli to regulate gene expression in bacteria.

## Introduction

Mechanical forces have long been recognized as key contributors to the growth and function of organisms. In *On Growth and Form*, D’Arcy Thompson highlighted the relationship between mechanical forces and the morphology and physiology of living organisms (1). In mammalian systems, mechanical forces regulate a wide variety of processes including cell differentiation during development (2, 3), disease initiation and progression (4), and tissue homeostasis (5). In tissues and organs with load-bearing functions, mechanical forces often act as the primary signal that initiates tissue remodeling and repair, thereby enabling the tissue to adapt to the mechanical challenges of the environment and quickly return to load bearing. Tissue remodeling thereby maintains homeostasis of mechanical function by balancing the removal of damaged tissue with tissue synthesis. Load-bearing structures including bone (6), blood vessels (7), and the plant cytoskeleton (8, 9) use mechanosensitive mechanisms to maintain mechanical function.

Most studies of mechanobiology focus on eukaryotic systems, although recent evidence has highlighted the importance of mechanical forces in prokaryotes. In bacteria, extracellular appendages including flagella and type IV pili extend from the cell body to sense and respond to mechanical cues in the environment. Flagellar motor unit assembly and disassembly respond to increases and decreases in external mechanical load (10, 11). Physical inhibition of flagellar rotation by contact with a surface generates reaction forces within the molecular motor, which stimulate surface adhesion and biofilm formation (12, 13). Type IV pili are motorized fibers that extend and retract to interact with the environment. In addition, mechanosensing by Type IV pili promotes biofilm formation (14) and the release of virulence factors (15), and it guides motility after collisions in the environment (16).

The cell envelope is the primary load-bearing component of bacteria and is also sensitive to mechanical forces. Stretch-activated ion channels within the cell membrane rapidly respond to changes in osmolarity by opening due to membrane stretching, leading to increased survival following hypo-osmotic shock (17). Mechanical stress and strain within the cell envelope also affect the assembly of trans-envelope efflux complexes; for example, assembly and function of the trans-envelope multicomponent efflux pump CusCBA is impaired by increases in octahedral shear stress within the cell envelope (18). Mechanical stress within the cell envelope also affects the locations of insertion of new cell wall in bacteria submitted to bending, with greater amounts of cell wall inserted at regions of greater tensile strain (19, 20). Although these mechanosensitive mechanisms within the cell envelope are well recognized, none of the mechanisms identified to date have been shown to regulate gene expression related to the remodeling of cell wall, an essential component of the cell envelope. If mechanosensitive mechanisms are involved in remodeling and homeostasis of the cell envelope, mechanical stress and strain would be expected to regulate the synthesis of components of the cell envelope.

*Vibrio cholerae*, the causative agent of cholera disease, survives rapid changes in osmolarity during the transition between fresh and brackish water, marine environments, and the intestines of a host. Osmolarity changes result in fluctuations in turgor that alter mechanical stress in the cell envelope. The primary structures counteracting turgor are the peptidoglycan (PG) cell wall and the outer membrane (OM) (21). To maintain adequate mechanical properties, bacteria must therefore maintain mechanical integrity of both PG and OM strength, and this indeed seems to be the case for the OM (21, 22). If and how PG strength is homeostatically controlled is poorly understood.

In *Vibrio cholerae*, the VxrAB two-component system is the major cell wall stress response system (23–25). VxrAB is induced by exposure to cell wall-acting antibiotics and overexpression of cell wall lytic enzymes like endopeptidase ShyA (25, 26). Upon induction, VxrAB upregulates its own expression, as well as cell wall synthesis functions, including the PG translocase MurJ, the major PG synthases (penicillin-binding proteins, PBPs), and PG precursor synthesis genes (Fig. 1A) (25). Consequently, VxrAB activation results in increased cell wall content and enhanced resistance to osmotic shock (25). These mechanisms are consistent with the idea that VxrAB contributes to cell wall homeostasis, similar to WalKR in the Gram-positive bacterium *Bacillus subtilis* (27). Consistent with this idea, a Δ*vxrAB* mutant exhibits increased cell width during normal growth (25), a finding expected if cell wall stiffness were impaired but turgor pressure remained the same. Importantly, VxrAB is also essential for survival after exposure to cell wall-acting antibiotics such as beta-lactams (25, 26). Beta-lactam exposure induces large-scale alterations of cell envelope mechanical properties, including a change from a rigid rod-shaped cell contained by a cell wall, to a membranous spheroplast (28, 29). Recovery from the spheroplast state relies primarily on VxrAB (26), presumably since rod shape regeneration requires increased cell wall synthesis. Thus, the VxrAB system modulates the mechanical properties of the cell in response to imbalances in PG turnover. The signal resulting in VxrAB induction, however, is still unknown.

**Figure 1.**
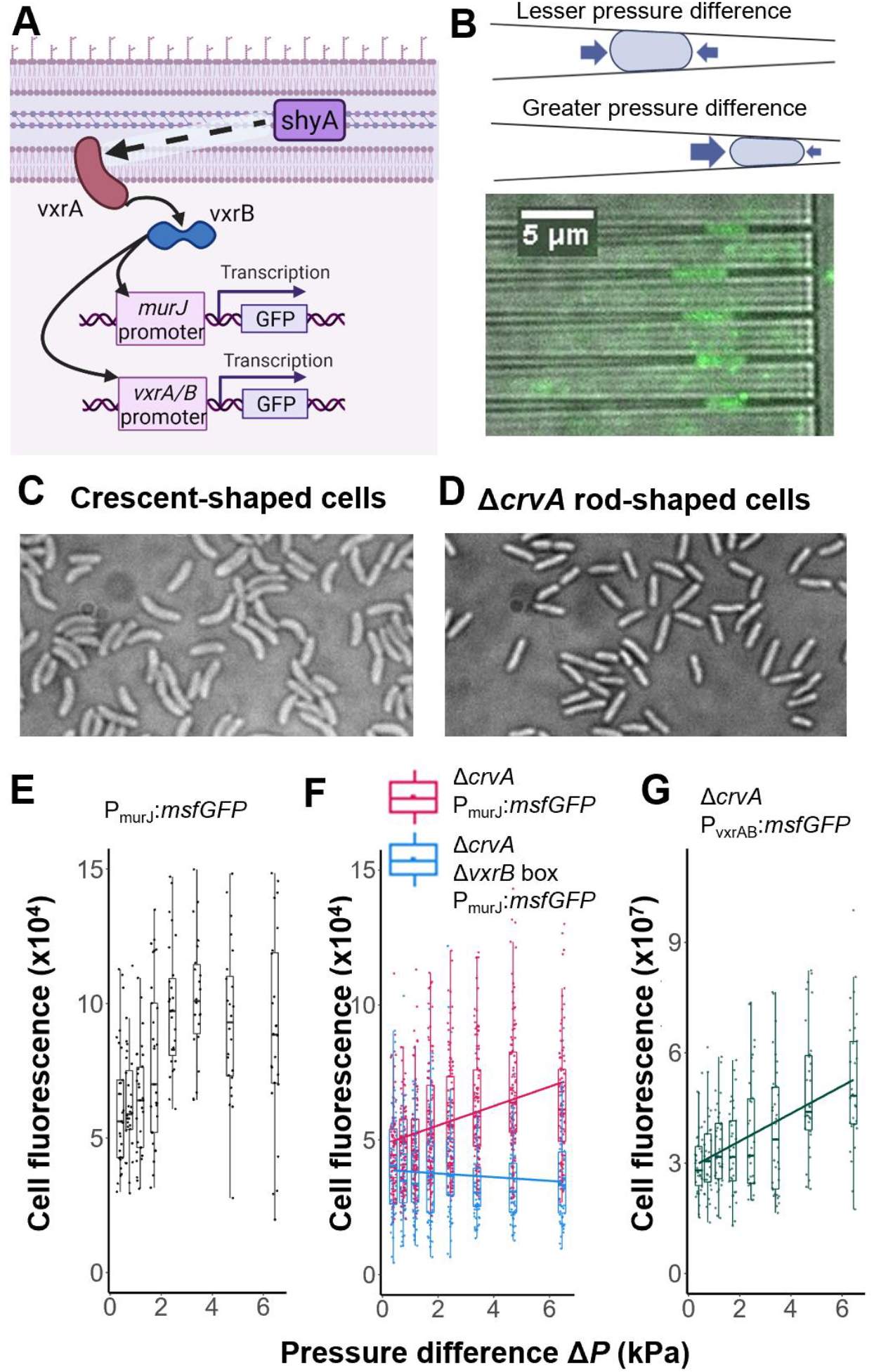
VxrAB signaling responds to mechanical stress from extrusion loading. (A) When activated, inner membrane histidine kinase VxrA phosphorylates response regulator VxrB, which regulates gene expression of regulons that include MurJ and VxrAB. Transcriptional msfGFP fusions for MurJ and VxrAB were used as reporters for VxrAB signaling. (B) Extrusion loading microfluidic device for applying mechanical loading. *Top*: Cells deform more and experience greater mechanical loading at a greater pressure difference. *Bottom*: Example transmission image with a fluorescent overlay of Δ*crvA* P_murJ_:*msfGFP* cells experiencing extrusion loading. (C) *Vibrio cholerae* are crescent-shaped. (D) *Vibrio cholerae* with *crvA* deletion are rod-shaped. (E) Single cell fluorescence of P_murJ_:*msfGFP* cells vs pressure difference. Solid line is a linear regression. (F) Single cell fluorescence of Δ*crvA* P_murJ_:*msfGFP* cells (pink) and Δ*crvA* Δ*vxrB* box P_murJ_:*msfGFP* cells (blue) vs pressure difference. Solid lines are linear regressions. Slope of Δ*crvA* P_murJ_:*msfGFP* cell (pink) is greater than the slope of the Δ*crvA* Δ*vxrB* box P_murJ_:*msfGFP* cells (blue) (P = 2e-12). (G) Single cell fluorescence of Δ*crvA* P_vxrAB_:*msfGFP* cells vs pressure difference. Solid line is a linear regression.

Here we study VxrAB controlled gene expression during mechanical stimulation of the cell envelope of *Vibrio cholerae*. We applied mechanical stress and strain to the bacterial cell envelope using three distinct loading modalities: extrusion loading within a microfluidic device, whole cell compression, and hydrostatic pressure. In each of these situations we find that bacteria experiencing greater magnitudes of mechanical loading exhibit greater VxrAB activation. Our findings support the idea that mechanical stress and strain play a role in regulating cell envelope synthesis and homeostasis and demonstrate the existence of gene regulatory systems that can be induced by mechanical stress in the cell envelope.

## Results and Discussion

### VxrAB signaling is activated by extrusion loading

To investigate the role of mechanical stress on VxrAB signaling, we used a custom microfluidic device to apply controlled, reproducible mechanical loads in a process we call “extrusion loading.” Extrusion loading uses fluid pressure to push bacteria into narrow tapered channels with sub-micron dimensions (Fig. 1B) (18, 30). Cells become lodged in the tapered channels and experience mechanical forces as they are deformed by the channel walls. The pressure difference (Δ*P*) across the tapered channel regulates the magnitude of mechanical stress experienced by a trapped cell. Cells submitted to greater pressure difference inside the tapered channels travel further into the tapered channels and experience greater deformation by the channel walls and greater magnitudes of mechanical stress. Analytical and finite element models indicate that extrusion loading results in increases in axial tensile stress, reductions in hoop tensile stress, and increases in octahedral shear (shape-changing) stress in the cell envelope (18).

We used a transcriptional P_murJ_:*msfGFP* fusion as a well-established reporter for VxrAB signaling (Fig. 1A). MurJ encodes for lipid II flippase for peptidoglycan precursor, and flipping the peptidoglycan precursor into the periplasmic space is critical for cell wall assembly (31). The response regulator VxrB has a high affinity for direct binding of the murJ promoter (26), and *murJ* expression is consequently strongly controlled by VxrAB (25), rendering this construct a robust readout of VxrAB activation. We applied extrusion loading to P_murJ_:*msfGFP* cells for two hours, then measured the fluorescence of individual cells to quantify the msfGFP expression under MurJ promoter control. The fluorescence of P_murJ_:*msfGFP* cells increased with increasing magnitude of extrusion loading (Fig. 1E), supporting the idea that VxrAB signaling is mechanosensitive.

*Vibrio cholerae* cells naturally exhibit a crescent shape (Fig. 1C). The straightening of a crescent-shaped cell inside the microfluidic device during extrusion loading results in additional mechanical stresses including greater tensile stresses on the concave side of the cell and compressive stresses on the convex side. To determine whether the mechanosensitive fluorescent response was solely due to the stresses caused by cell straightening, we created rod-shaped *V. cholerae* by deleting *crvA* (Fig. 1D) (32) and submitted Δ*crvA* P_murJ_:*msfGFP* cells to extrusion loading. The fluorescence of Δ*crvA* P_murJ_:*msfGFP* cells also increased with increasing pressure difference (Fig. 1F, magenta), suggesting that cell curvature is not necessary for the mechanosensitive fluorescent response of P_murJ_:*msfGFP* cells during extrusion loading. The difference in slope of the fluorescence vs pressure difference between the crescent-shaped and the rod-shaped cells may be explained by the additional stresses that crescent-shaped cells experience from straightening. All subsequent experiments were performed with rod-shaped, Δ*crvA* cells. The autofluorescence signal of Δ*crvA* non-GFP producing cells was not increased at greater magnitudes of extrusion loading (*SI Appendix*, Fig. S5).

To confirm that the mechanosensitive response was due to VxrB-activated expression of P_murJ_:*msfGFP*, we used a mutant with a partial deletion of the VxrB binding site on the MurJ promoter 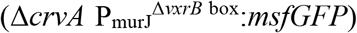; we previously established that this mutant indeed lacks VxrAB-responsiveness (26). Under extrusion loading, the fluorescence of such mutant cells did not vary with pressure difference (Fig. 1F, blue), demonstrating that VxrB activation is required for the mechanosensitive increase in P_murJ_:*msfGFP* expression. In addition to its role in MurJ induction, VxrAB signaling is also autoregulated (Fig. 1A) (26). To confirm that the mechanosensitive response of VxrAB was not limited to the MurJ promoter, we also used a P_*vxrAB*_*:msfGFP* reporter strain as an alternative readout of VxrAB activation. Cell fluorescence of P_*vxrAB*_*:msfGFP* cells increased at greater pressure difference (Fig. 1G), confirming that mechanical stress in the cell envelope regulates VxrAB signaling. We conclude that VxrAB activation is sensitive to mechanical stress from extrusion loading.

### VxrAB signaling is activated by other forms of mechanical loading: compression and hydrostatic pressure

To confirm that the mechanosensitive response of VxrAB signaling is not specific to the microfluidic device environment or the forms of mechanical stresses generated by extrusion loading, we investigated the response of VxrAB signaling to two other methods of mechanical loading: compression and hydrostatic pressure.

To apply compression, cells were sandwiched between agarose gel and a weighted glass slide (Fig. 2A). At greater magnitudes of applied force, cells experienced greater mechanical loading and deformed to a greater cell width in the plane of imaging (*SI Appendix*, Fig. S4). Compression causes increases in tensile stress in the axial and hoop directions and overall increase in octahedral shear (shape-changing) stress. P_murJ_:*msfGFP* cells were submitted to compression loading for two hours (the same duration that cells experienced under extrusion loading), and the fluorescence of individual cells was then measured to quantify expression of P_murJ_:*msfGFP*. Cell fluorescence increased 30% with increasing magnitude of applied compression force (Fig. 2B, left), indicating that VxrAB signaling is sensitive to compression loading. In contrast, upon deleting the *vxrB* box in the promoter (Δ*vxrB* box), cell fluorescence did not vary significantly with applied compression force (Fig. 2B, middle), demonstrating that the mechanosensitive increase in cell fluorescence for P_murJ_:*msfGFP* cells under compression also required VxrB activation. We again used P_*vxrAB*_*:msfGFP* cells as a secondary reporter for VxrAB signaling. Cell fluorescence of P_*vxrAB*_*:msfGFP* cells increased with increasing applied compression force (Fig. 2B, right), further supporting the idea that the mechanosensitive response to compression is mediated by VxrAB. We note that msfGFP fluorescence shows larger variance among individual cells during compression loading than during extrusion loading, which is likely because compression does not apply as rigorously controlled cell contact and load distribution.

**Figure 2.**
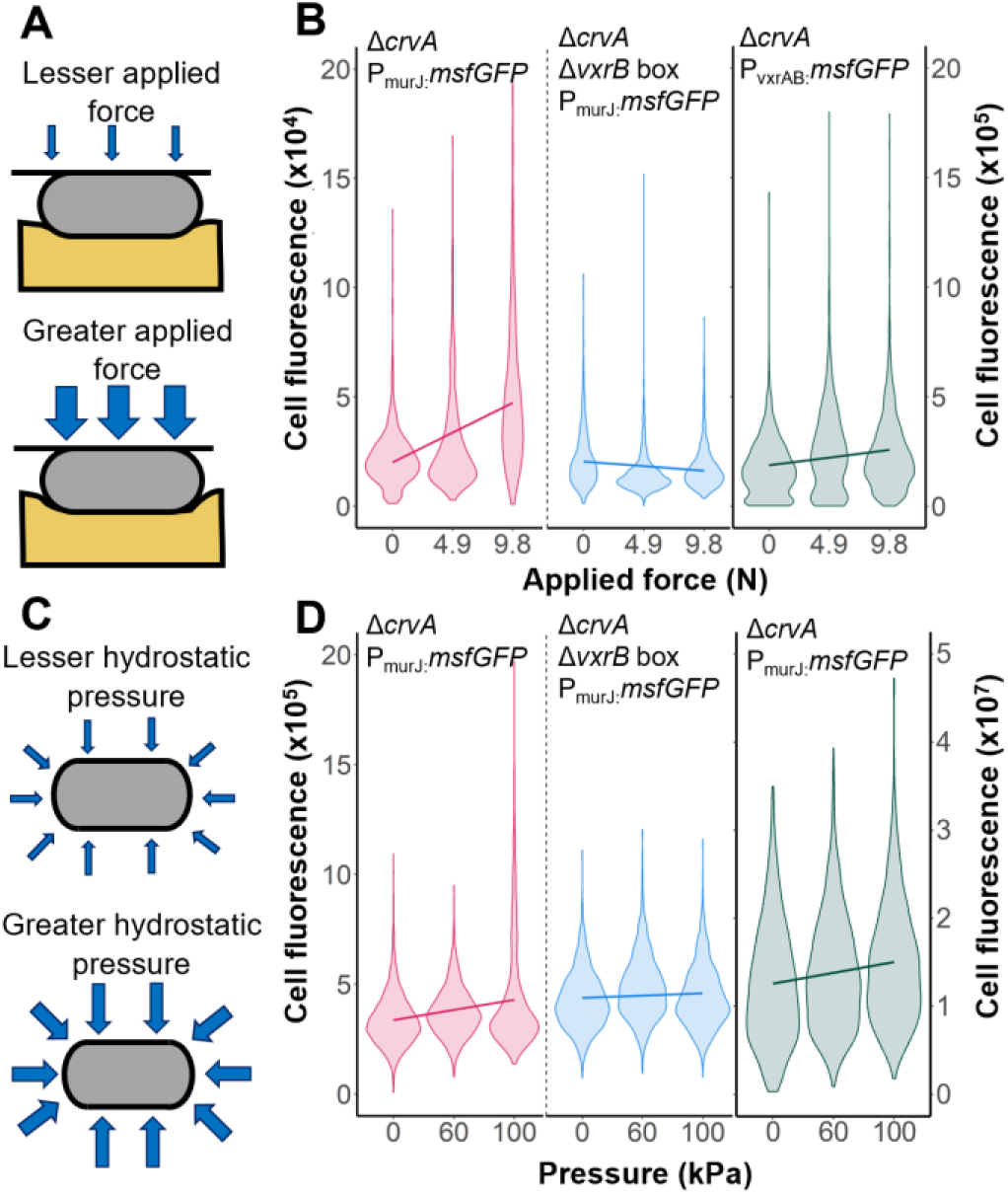
VxrAB signaling responds to mechanical stress from compression and hydrostatic pressure. (A) During compression, cells are sandwiched between agarose gel and a weighted glass slide. (B) Single cell fluorescence of Δ*crvA* P_murJ_:*msfGFP* cells (pink), Δ*crvA* Δ*vxrB* box P_murJ_:*msfGFP* cells (blue), and Δ*crvA* P_vxrAB_:*msfGFP* cells (green) vs applied compression force. Solid lines are linear regressions. The slope of Δ*crvA* P_murJ_:*msfGFP* cells (pink) is greater than the slope of the Δ*crvA* Δ*vxrB* box P_murJ_:*msfGFP* cells (blue) (P < 2e-16). (C) Hydrostatic pressure applies force equally to all surfaces of the cell. (D) Single cell fluorescence of Δ*crvA* P_murJ_:*msfGFP* cells (pink), Δ*crvA* Δ*vxrB* box P_murJ_:*msfGFP* cells (blue), and Δ*crvA* P_vxrAB_:*msfGFP* cells (green) vs hydrostatic pressure. Solid lines are linear regressions. The slope of Δ*crvA* P_murJ_:*msfGFP* cells (pink) is greater than the slope of the Δ*crvA* Δ*vxrB* box P_murJ_:*msfGFP* cells (blue) (P < 2e-16).

Next we assessed the role of hydrostatic pressure in VxrAB induction. Hydrostatic pressure exerts force perpendicular to all surfaces of the cell (Fig. 2C). Extreme hydrostatic pressure (>50 MPa) causes changes to RNA synthesis and DNA replication and can cause cell death (33); here we apply mild hydrostatic pressure of 0-100 kPa (for reference, a bacterium 10 meters underwater experiences approximately 100 kPa hydrostatic pressure). Hydrostatic pressure causes reductions in tensile stress in the axial and hoop directions and overall increases in octahedral shear (shape-changing) stress. MurJ was again used as a reporter for VxrAB signaling. P_murJ_:*msfGFP* cells submitted to two hours of hydrostatic pressure. Cell fluorescence increased 28% with increasing magnitude of applied compression force (Fig. 2D, left), demonstrating that VxrAB signaling is also sensitive to hydrostatic pressure. Upon deleting the *vxrB* box in the promoter, cell fluorescence did not vary with applied hydrostatic pressure (Fig. 2D, middle), confirming that the mechanosensitive increase in cell fluorescence for P_murJ_:*msfGFP* cells required VxrB activation. As in the experiments above, the P_vxrAB_:*msfGFP* reporter strain recapitulated the observations with P_murJ_:*msfGFP* (Fig. 2D, right). We conclude that VxrAB signaling is responsive to diverse methods of mechanical load application to the cell envelope.

### Cell wall turnover affects mechanosensitivity of VxrAB signaling

Our above results show that VxrAB signaling responds to mechanical stimuli, but it is unknown if the kinase VxrA is itself a mechanosensor or if VxrA activation is an indirect response to other changes caused by mechanical stress in the cell envelope. VxrAB is known to respond to cell wall damage caused by cell wall targeting antibiotics and overexpression of the endopeptidase ShyA (Fig. 1A) (25). Cell wall damage could trigger VxrAB mechanically by reducing cell envelope stiffness and thereby increasing deformation of the cell envelope caused by turgor, or VxrAB could respond to a chemical byproduct of cell wall degradation. Endopeptidases cleave peptide strands in the peptidoglycan, allowing for the insertion of new cell wall material and cell elongation, but also contribute to cell wall degradation after antibiotic exposure (34, 35). The endopeptidase ShyA plays a key role in cell wall homeostasis and cell elongation through controlled cell wall degradation in *V. cholerae* (36). ShyA overexpression, by increasing degradation of the cell wall, may cause reductions in cell envelope mechanical properties, resulting in greater mechanical deformation of the cell envelope under turgor pressure and a resulting increase in VxrAB signaling. Alternatively, ShyA-mediated degradation of the cell wall could cause the release of cell wall fragments or byproducts leading to the activation of VxrA. To explore the possibility that VxrAB mechanosensitive signaling is related to cell wall damage or degradation, we exposed mutants with ShyA deletion or ShyA overexpression to extrusion loading and measured the resulting changes in VxrAB signaling.

Under extrusion loading, the fluorescence of cells with ShyA deleted (Δ*shyA* P_murJ_:*msfGFP*) ceased to be correlated with pressure difference (Fig. 3A). When ShyA was reintroduced in trans, the mechanosensitive response was rescued: cell fluorescence increases with increasing pressure difference during extrusion loading for Δ*shyA* ShyA++ P_murJ_:*msfGFP* (Fig. 3B). However, we noticed that upon deleting ShyA, cells did not travel as far into the tapered channels during extrusion loading as cells with normal ShyA expression (Fig. 3C), potentially suggesting that cells defective in major EP activity experienced insufficient cell deformation compared with cells with normal ShyA expression. However, cells with ShyA deleted outside of the microfluidic devices were wider than cells with normal ShyA expression (Fig. 3D, top vs. bottom). Due to this width increase, Δ*shyA* P_murJ_:*msfGFP* cells might experience greater mechanical strain/deformation than the P_murJ_:*msfGFP* cells despite not traveling as far in the tapered channel. Indeed, when comparing cell width inside and outside the tapered channels, Δ*shyA* P_murJ_:*msfGFP* cells experienced a greater percentage decrease (average of 40% decrease in cell width) in cell width as compared to P_murJ_:*msfGFP* cells (average of 25% decrease in cell width). Therefore, the lack of mechanosensitive response in the Δ*shyA* strain is likely not due to insufficient deformation of the cell envelope. Furthermore, the cells in tapered channels for both groups experience the same pressure difference, so the loss of mechanosensitivity in the Δ*shyA* P_murJ_:*msfGFP* cells is not due to differences in applied mechanical force. Since VxrAB baseline levels were elevated in the Δ*shyA* mutants (indicating functional VxrA), the inability of VxrAB to respond to mechanical loading indicates that it is unlikely that VxrAB directly senses mechanical stress and strain and that mechanosensation of VxrAB is either secondary to ShyA activity or indirectly due to the deletion of ShyA (perhaps related to consequent changes associated with increased expression of other endopeptidases or perturbations cell wall mechanical properties or homeostasis in the absence of ShyA). Indeed, *ΔshyA* P_murJ_:*msfGFP* cells exhibit noticeably perturbed physiology with blebs, curves, and abnormal shape, suggesting the possibility of impaired cell envelope mechanical properties that might reduce cell stiffness (Fig. 3E). We speculate that VxrAB signaling in response to mechanical stress is related to cell wall fragments released during mechanical loading. Mechanical stress might enhance ShyA activity through substrate availability, thus resulting in production of more PG fragments that are sensed by VxrA. Indeed, PG lytic enzymes have been previously hypothesized to respond to PG stretching state (37). Another possibility is that the absence of ShyA results in changes in PG structure in a way that makes it non-conducive to VxrAB-mediated mechanosensing. Although the mechanism of VxrAB mechanosensation remains unclear, our findings suggest that cell envelope remodeling and homeostasis are at least in part mechanically regulated.

**Figure 3.**
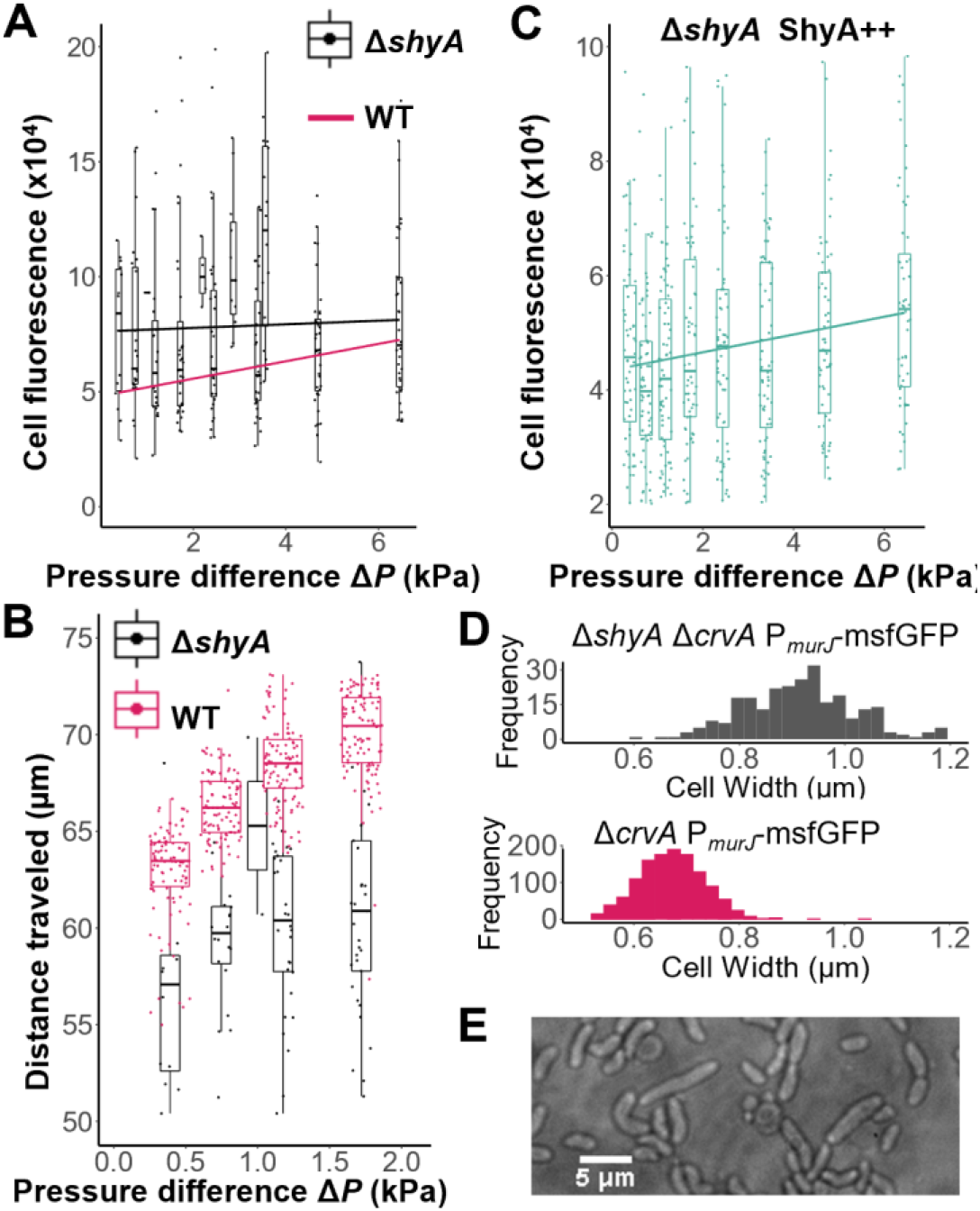
ShyA and VxrAB mechanosensitivity. (A) Single cell fluorescence of Δ*shyA* Δ*crvA* P_vxrAB_:*msfGFP* cells vs pressure difference in extrusion loading (black). Δ*crvA* P_murJ_:*msfGFP* linear fit trendline shown in pink from Fig. 1F. Solid black line is a linear regression for the Δ*shyA* Δ*crvA* P_vxrAB_:*msfGFP* cells. The slope of Δ*crvA* P_vxrAB_:*msfGFP* cells (pink) is greater than the slope of the Δ*shyA* Δ*crvA* P_vxrAB_:*msfGFP* cell (P = 1e-7). (B) Single cell fluorescence of Δ*shyA* Δ*crvA* ShyA++ P_murJ_:*msfGFP* cells vs pressure difference. Solid green line is a linear regression. (C) Distance traveled in the tapered channel for Δ*shyA* Δ*crvA* P_murJ_:*msfGFP* cells (black) and Δ*crvA* P_murJ_:*msfGFP* cells (pink) vs pressure difference. (D) *Top*: Δ*shyA* Δ*crvA* P_murJ_:*msfGFP* cell width in a petri dish. *Bottom*: Δ*crvA* P_murJ_:*msfGFP* cell width in a petri dish. (E) Example image of Δ*shyA* Δ*crvA* P_murJ_:*msfGFP* cell morphology grown in M9 medium in a petri dish.

## Conclusions

We have demonstrated that the two-component signaling system VxrAB is activated by mechanical stress in the cell envelope caused by three distinct mechanical loading modalities: extrusion loading, compression, and hydrostatic pressure. These findings indicate that mechanosensitive mechanisms within the cell envelope can regulate gene expression involved in cell wall remodeling. While VxrAB responds to mechanical stress, it is unlikely that physical forces directly regulate VxrAB function suggesting that the primary effect of physical forces occurs on another process that remains to be identified.

Although our findings suggest that VxrA does not respond directly to mechanical stress and strain, our findings support the idea that mechanical stress and strain may play a role in controlling cell wall homeostasis. The ability of mechanical stress and strain to contribute to the remodeling of a load-bearing component such as the cell wall is a powerful means of maintaining the function of the cell envelope. While we are unaware of other studies directly investigating the effects of mechanical stress on cell wall maintenance in other bacteria, there are other two-component systems that regulate cell wall remodeling. The two component system WalKR in *Bacillus subtilis* responds to degradation of cell wall constituents to regulate cell wall remodeling (27). Additionally, recent findings indicate that the outer membrane is also capable of carrying mechanical loads (21), opening the possibility that the synthesis and transport of outer membrane components may be sensitive to mechanical stress.

The mechanosensitive nature of VxrAB suggests potential applications in the field of synthetic biology. Gene regulatory mechanisms that respond to the external environment are key tools in the field of synthetic biology. Systems that respond to target chemicals, light, temperature, and pH have been used to control synthetic gene circuits (38). Our work demonstrates that two-component systems can respond to mechanical stress in the cell envelope, providing an additional mechanical mechanism for stimulating gene circuits for synthetic biology applications.

## Contributions

C.E.H.: wrote the manuscript, performed mechanical stimuli experiments (extrusion loading, hydrostatic pressure, and compression) and imaging, analyzed and curated data, performed statistical analysis, fabricated microfluidic devices, wrote script for image processing, performed image processing; W.Z.: wrote the manuscript, performed mechanical stimuli experiments (extrusion loading, hydrostatic pressure, and compression) and imaging, analyzed data, performed image processing; J.-H.S.: constructed strains; E.v.W.: performed compression experiments, performed image processing; E.C.: performed image processing; J.L. performed hydrostatic pressure experiments; Z.W.: contributed to sample preparation and imaging; T.D. designed and directed research, edited manuscript; P.C. designed and directed research, edited manuscript; C.J.H. designed and directed research, edited manuscript

## Acknowledgements

NSF 2055214, 2135586 (C.J.H.), NSF GRFP DGE-1650441 (C.E.H.). This work was performed in part at the Cornell NanoScale Facility, a member of the National Nanotechnology Coordinated Infrastructure (NNCI), which is supported by the National Science Foundation (Grant NNCI-2025233). Research on VxrAB in the Dörr lab is supported by National Institutes of Health (NIH/NIAID) grant R01AI143704.

